# Mosquito invasion via the global shipping network is slowed in high-risk areas by on-shore and ship-board monitoring

**DOI:** 10.1101/2022.08.29.505734

**Authors:** Janna R. Willoughby, Benjamin A. McKenzie, Jordan Ahn, Todd D. Steury, Christopher A. Lepzcyk, Sarah Zohdy

## Abstract

The global shipping network (GSN) has been suggested as a pathway for the establishment and reintroduction of *Aedes aegypti* and *Aedes albopictus* primarily via the tire trade. We used historical maritime movement data in combination with an agent-based model to understand invasion risk in the United States Gulf Coast and how the risk of these invasions could be reduced. We found a strong correlation between the total number of cargo ship arrivals at each port and likelihood of arrival by both *Ae. aegypti* and *Ae. albopictus*. Additionally, in 2012, 99.2% of the arrivals into target ports had most recently visited ports occupied by both *Ae. aegypti* and *Ae. albopictus*, increasing risk of *Aedes* invasion. Model results indicated that detection and removal of mosquitoes from containers when they are unloaded at a port may be more effective in reducing the establishment of mosquito populations compared to eradication efforts that occur while onboard the vessel, suggesting detection efforts should be focused on unloaded containers. To reduce the risk of invasion and reintroduction of *Ae. aegypti* and *Ae. albopictus*, surveillance and control efforts should be employed when containers leave high risk locations and when they arrive in ports at high risk of establishment.

## Introduction

The globalization of trade and travel has allowed many invasive species to disperse and establish themselves in novel locations and at distances much farther than their natural dispersal abilities should allow [1, 2]. These dispersal events are fueled by our increasingly interconnected world [1]. The global shipping network (GSN), which currently accounts for >80% of international trade, has expanded dramatically in the past 50 years and is expected to increase by at least 240% by 2050 [2, 3]. Notably, the GSN acts as a significant pathway for the long-distance transport of invasive and other non-native organisms to novel locations [4, 5]. Non-native aquatic species are often transported in the ballast water or attached to the hulls of vessels [6, 7], while terrestrial species are often accidentally transported with the cargo [2, 8].

International maritime trade and the GSN have been instrumental in the global invasions of several medically important *Aedes* spp. mosquitoes, most notably *Aedes* (Stegomyia) *aegypti* (L.) and *Aedes* (Stegomyia) *albopictus* (Skuse) [9–11] that are the primary vectors of globally significant arboviruses including the dengue fever, chikungunya, Zika, and yellow fever viruses [12–15]. Although both species have relatively short flight ranges and therefore limited in self-powered dispersal [16], these species have adaptations which have allowed them to use the GSN for dispersal and establish near-global distributions [17, 18]. For example, the expansion of the *Ae. aegypti* range was marked by a shift in blood-feeding behavior [19]: anthropophagy in *Ae. aegypti* is believed to have developed during long ship crossings in the pre-industrial era, where selection against zoophagy would have been strong [10]. In addition, *Ae. aegypti* and *Ae. albopictus* can oviposit in artificial containers allowing them to thrive in highly urban environments [20, 21]. Adaptations to anthropogenic environments, combined with the unique ability of their eggs to survive desiccation for extended periods [20, 22], have allowed these mosquitoes to be transported globally through the GSN among potted plants and used tires [17, 23, 24].

While climate change alters global patterns of habitat suitability for both *Ae. aegypti* and *Ae. albopictus* [25, 26], both species continue to expand in most parts of the world thanks to new maritime introduction events and overland spread [19]. Because of widespread reductions in vector control efforts towards the end of the 20^th^ century and continuous reintroductions, *Ae. aegypti* has also reestablished itself in parts of its range from which it was once extirpated, including parts of the southeastern US [27, 28]. The most effective strategy in limiting the spread of invasive species, including *Ae. aegypti* and *Ae. albopictus*, is the implementation of effective biosecurity measures at points of entry [29] using early detection and rapid response to prevent establishment [30]. Because resources available for effective early detection and rapid response networks are generally limited, the identification of high risk locations for the importation and reintroduction of invasive species is critical for effective biosecurity [8, 29, 30].

The Gulf Coast of the United States (Figure 1) has been identified as a region at risk for the emergence and establishment of Zika virus and other arboviruses associated with *Aedes* spp. mosquitoes due to its warm and humid climate as well as the presence of many key transportation hubs (airports and seaports) within the region [31, 32]. However, while there have been outbreaks of Zika, dengue, and chikungunya in the U.S. Gulf States, these outbreaks do not compare in magnitude with those experienced in nearby Latin America [33–36]. This discrepancy may be partially explained by varying vector control and public health efforts as well as differences in housing style and lifestyle between affected countries, but is likely also partly due to differences in vector competence between mosquito populations [37–40]. Variation in vector competence between populations can occur at relatively fine spatial scales [41], meaning that continuous reintroduction of vectors into a region can influence local mosquito competency for various viruses. Thus, halting the genetic flow between disparate mosquito populations will aid in preventing the establishment of arboviral diseases in new locations. Developing models that predict reinvasions by *Ae. aegypti* and *Ae. albopictus* and identify the best strategies for targeted biosurveillance and vector control in ports could help to alert public health officials to potential threats and support optimized biosecurity efforts to halt reinvasion.

**Figure 1.**
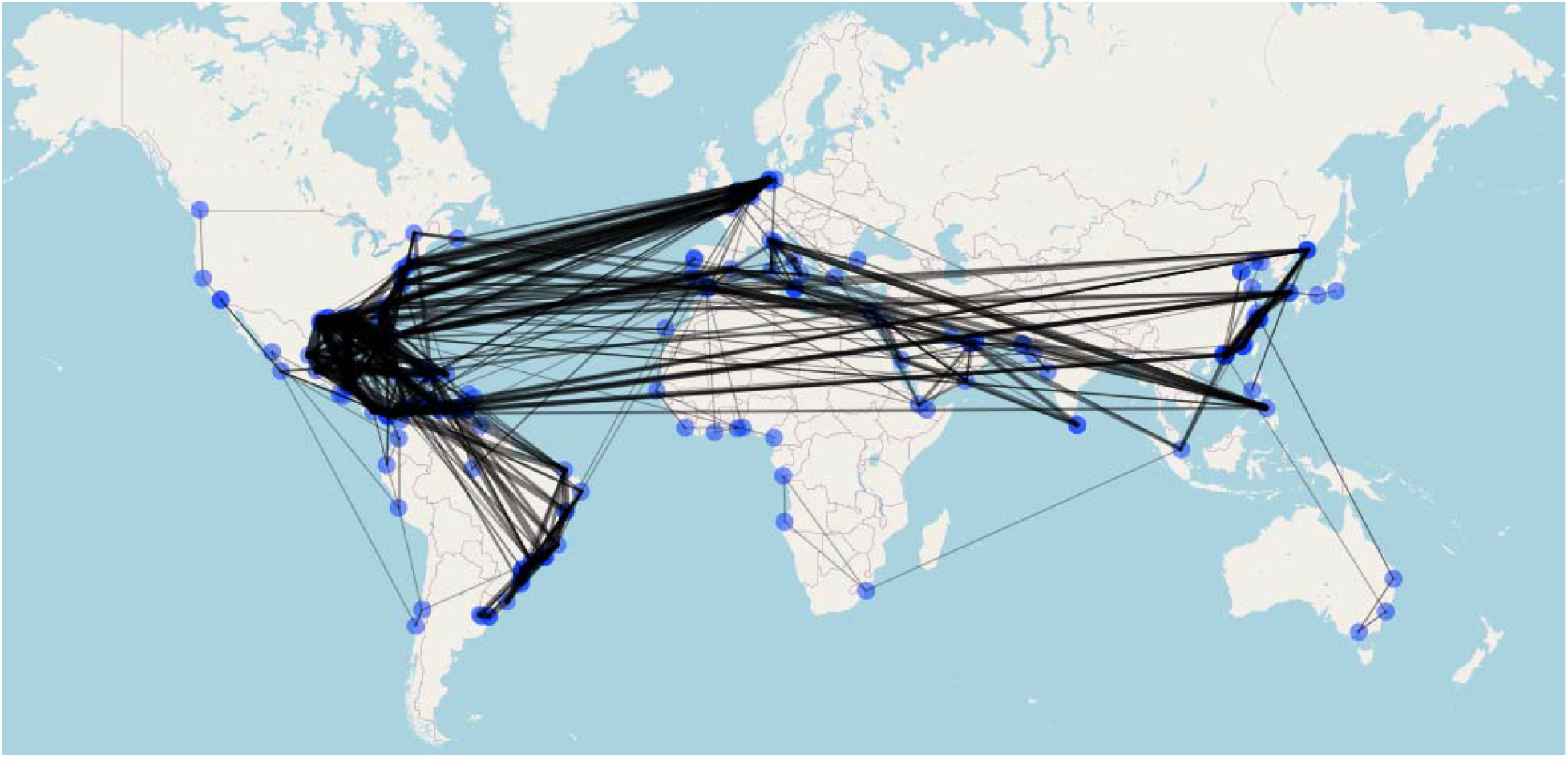
The Global Shipping Network containing Gulf State ports included 213 ports located in 69 countries. Lines are logarithmically weighted to demonstrate connectivity between ports and indicate a highly connected shipping network.

The goal of this study was to first integrate available *Aedes* species distribution data and maritime movement data to identify ports at high risk for importation of *Ae. albopictus* and *Ae. aegypti* via the GSN. Following this, we sought to understand how invasion risk could be minimized at high-risk ports and ports connected to these high-risk areas using biosurveillance. To do this, we used an agent-based model to quantify how port detection and removal protocols could be used to limit the establishment of mosquito vectors in new areas. This work may help officials concentrate biosecurity efforts to prevent further mosquito invasion and potential importation of vector-borne pathogens.

## Results

We used data detailing every fully cellular container ship that arrived into major US ports on the Gulf of Mexico between January 1^st^ and December 31^st^, 2012. These data were recorded by automatic identification system transponders, which are installed on every large ship and at every port and canal in world and automatically report data on ship size, location. These data included the previous ten ports of call for each ship before arriving in one of seven US ports (Figure 1).

We documented 1,921 arrivals and departures of 204 container ships. Using *Aedes* distributions maps [18], we determined that only 39 (18.3%) of the 213 ports within our network (distributed across 69 countries) were free of *Ae. aegypti* and *Ae. albopictus* populations; 140 (65.7%) ports within our network hosted populations of *Ae. aegypti*, 148 (69.4%) hosted populations of *Ae. albopictus*, and 114 (53.5%) ports within our network hosted populations of both *Ae. aegypti* and *Ae. albopictus* (Supplementary Figure 1).

### Invasion risk assessment

We used pathway-based, first-order Markov models to determine which ports along the US Gulf Coast were at the highest risk for importation of *Ae. aegypti* and *Ae. albopictus* along with container cargo shipped via maritime trade routes during the year examined, 2012. Given that a ship loads and unloads cargo with each stop, our models also assume that some potential exists for infestation of the ship by mosquitoes at each stop at a port occupied by these species; because there is also some probability of cargo containing the mosquitoes to be unloaded at each port, we considered all points on a route together to fully understand invasion patterns. We determined that port traffic is a strong indicator of probability of invasion; we found a strong correlation between the total number of cargo ship arrivals at each port and likelihood of arrival by *Ae. aegypti* (r^2^ = 0.999 P > 0.0001) and *Ae. albopictus* (r^2^ = 0.999, P > 0.0001) (Table 1). Of the 1,921 arrivals into target ports, 99.2% of ships (n=1,905) were moving from ports occupied by both *Ae. aegypti* and *Ae. albopictus* populations and only one arrival was coming from a port where neither species is commonly found (Supplementary Table 1). We also observed a high level of connectivity between ports (Supplementary Table 2). Combined, these data suggest high probability of invasion potential, including movement of mosquitoes from a previously invaded location to a new or other previously invaded location.

**Table 1.**
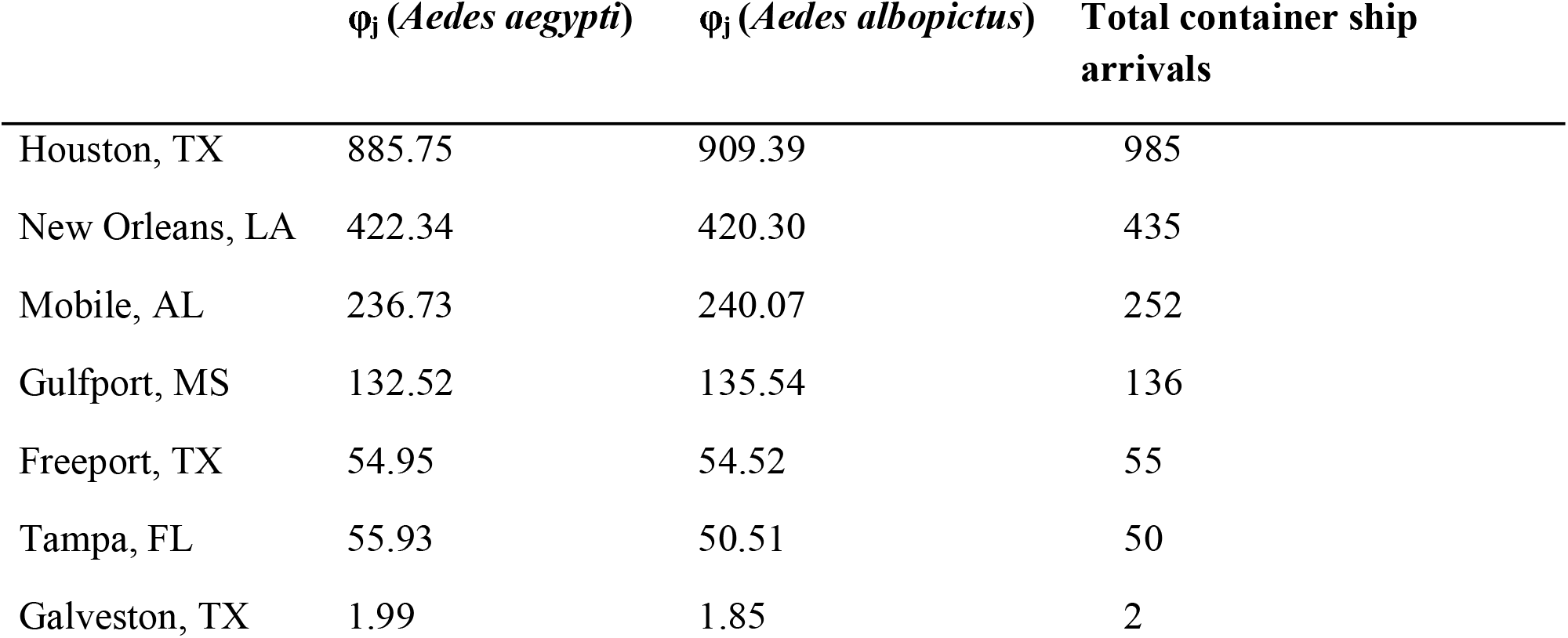
Ports along the Gulf Coast of the US with the highest relative likelihood of arrival (φ_*j*_) by *Ae. aegypti* and *Ae. albopictus* via the international maritime trade network given a constant transmission potential (λ) of 0.5. The total number of arrivals of fully cellular container ships at each port was strongly correlated with relative likelihood of arrival by both *Ae. aegypti* (r^2^ = 0.999, P > 0.0001) and *Ae. albopictus* (r^2^ = 0.999, P > 0.0001) during this time frame.

| | $\phi_j$ (*Aedes aegypti*) | $\phi_j$ (*Aedes albopictus*) | Total container ship arrivals |
| --- | --- | --- | --- |
| Houston, TX | 885.75 | 909.39 | 985 |
| New Orleans, LA | 422.34 | 420.30 | 435 |
| Mobile, AL | 236.73 | 240.07 | 252 |
| Gulfport, MS | 132.52 | 135.54 | 136 |
| Freeport, TX | 54.95 | 54.52 | 55 |
| Tampa, FL | 55.93 | 50.51 | 50 |
| Galveston, TX | 1.99 | 1.85 | 2 |

### Invasion risk mitigation

We designed an agent-based model to understand how to effectively mitigate mosquito invasion risk. In the model, shipping containers aboard maritime ships were treated as agents, where mosquitoes found in these vessels could be moved between ports. Each container was assumed to start its journey with enough mosquitoes that mosquito establishment at new ports was theoretically possible. Containers moved between ports and could be moved to shore at any port with varying probabilities; when containers were not moved to shore, containers could instead be transferred to a different cargo ship or remain on the original ship.

To combat the establishment of new mosquito vector populations at these simulated ports, we enacted port procedures for detecting and removing mosquitoes from shipping containers.

Containers were checked upon arrival to shore or transfer between ships, with a probability of removing mosquitoes ranging from 0 to 100% for each event. We found that detection and removal of mosquito infestations after unloading at the destination reduced the probability of mosquito establishment, but that detection and removal of mosquitoes after containers were transferred between vessels had little effect on mosquito population establishment because mosquito infestation in containers that are never moved to shore are somewhat irrelevant to invasion probability (Figure 2). Thus, on-shore mosquito detection and eradication efforts in maritime goods is more likely to prevent new invasions compared to on-ship detection and removal efforts.

**Figure 2.**
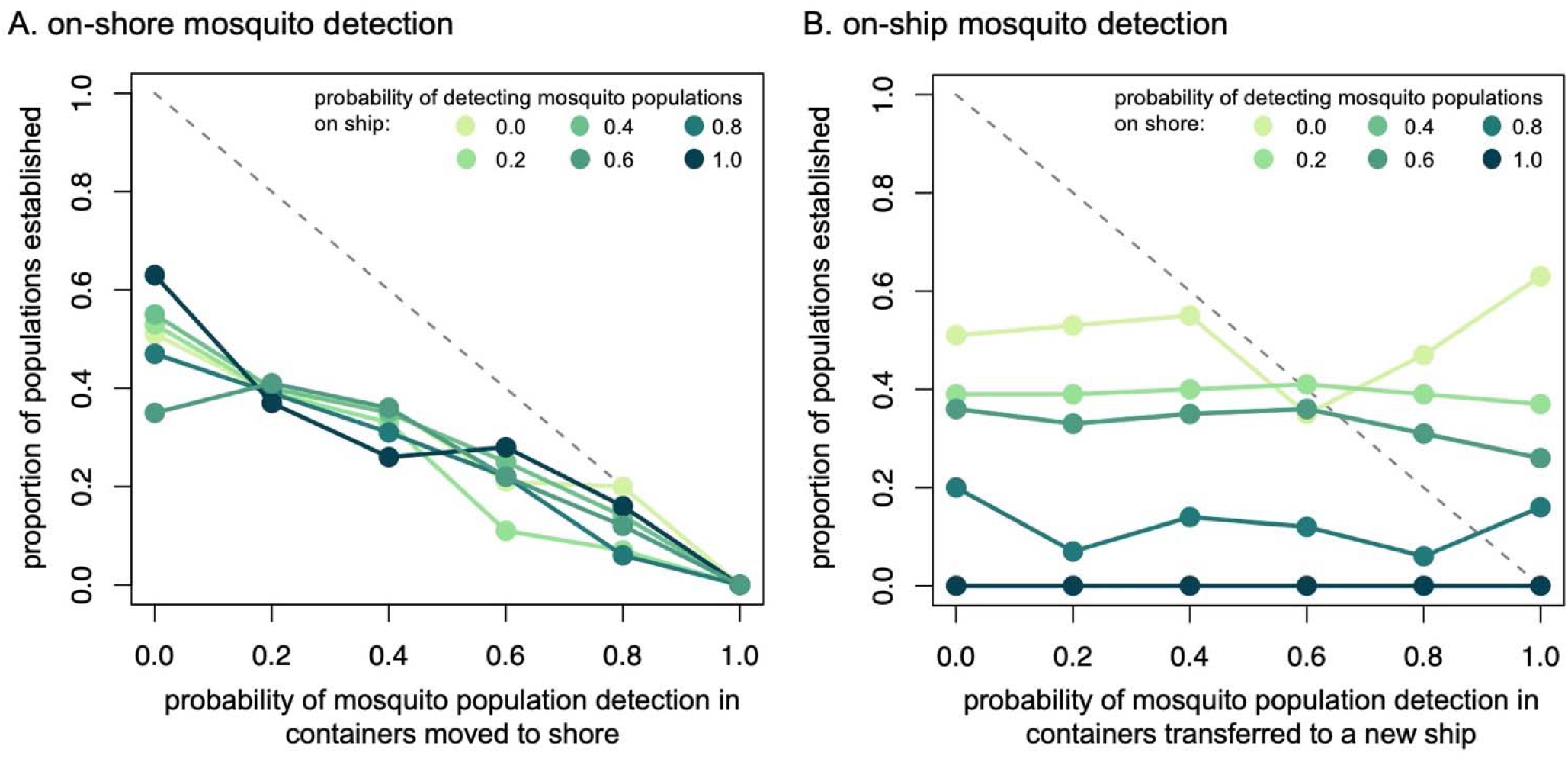
Effectiveness of on-shore and on-ship based mosquito detection and removal programs. A) Probability that mosquito-infected containers were detected in on-shore efforts and the percent of mosquito populations that established in new areas, illustrated across various on-ship efforts to detect container infections that also occurred. B) The percent of established mosquito populations given the probability that mosquito-infected containers were detected in ship-board efforts, displayed across land-based efforts levels that co-occurred. Across levels of effort and effectiveness, on-shore detection and eradication methods are more effective at preventing new mosquito infestations compared to ship-based efforts.

We also compared the number of stops a ship made, equivalent to the number of visited ports. In our model and in reality, the number of stops a ship makes controls the number of opportunities for transfer between ships and land and, therefore, the opportunities for mosquito detection, eradication, and establishment. We found that the number of stops did not interact with the effect of removing mosquitoes from unloaded containers (Figure 3a). However, there may be a weak interaction between number of stops and the effect of removing mosquitoes after ship-to-ship container transfer (Figure 3b); when the number of stops was small (e.g. 1-5), the proportion of established populations was more variable when compared within detection probabilities, but these differences decreased as the number of stops increased towards 10. This is because the repeated searching and removing of infestations meant that even if the probability of detecting the mosquito infestations in one search was small, with enough searches (i.e., stops) these infestations could be eventually detected and removed. This suggests that repeated searching on-board ships provides some additional protections from invasion even when detection probabilities are small.

**Figure 3.**
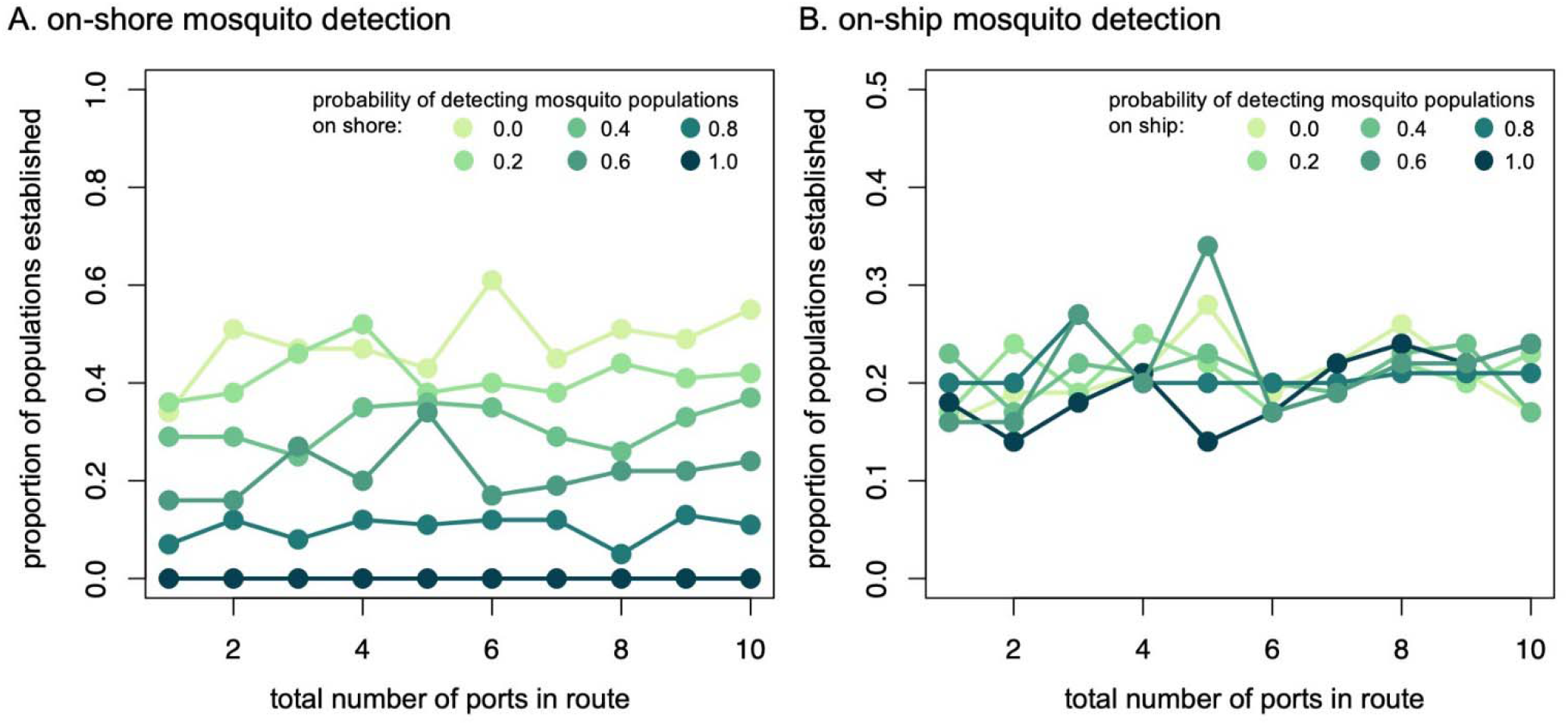
Comparison of the effect of number of ship ports (stops) in a route to probability of mosquito population establishment. In A, the colors show increasing effectiveness of mosquito removal from unloaded containers, when on-ship probability of detecting and removing was held constant at 60%. In B, the colors indicate effectiveness of removal from the transferred containers when on-shore probability of removal was held at 60%. Overall, the variation in probability of mosquito population establishment decreased with respect to ship-based detections, when the number of ports increased.

## Discussion

In our models, connectivity as measured by frequency of ship arrivals and previous ports of call, predicted likelihood of mosquito invasion and this has important ramifications for eventual invasion by *Aedes* species. For example, the Port of Houston, Texas represented by far the greatest risk for the invasion of *Ae. aegypti* and *Ae. albopictus* to other US ports along the coast of the Gulf of Mexico. In our 2012 dataset, the Port of Houston received more than double the arrivals of container ships as did the U.S. Gulf port with the next most arrivals, New Orleans. In fact, Houston received more arrivals during this period than did all six other major U.S. ports in the Gulf combined (Table 1). While more than three-quarters of container ships arriving in Houston had most recently come from a port outside the Gulf which hosted both *Ae. aegypti* and *Ae. albopictus*, the majority of traffic into other Gulf ports was internal, with arrivals coming from other Gulf ports (Supplementary Tables 1 and 2). These data are in line with other historical data on frequency of container ship arrivals and cargo tonnage, which show that Houston received more arrivals and handled more tonnage than any other port in the Gulf from 2016-2018, and that Houston handled a higher proportion of foreign arrivals and freight than did other Gulf ports [42].

While the total number of arrivals by container ships may not always indicate the highest likelihood of arrival by invasive species generally, the distributions and common occurrence of *Ae. aegypti* and *Ae. albopictus* within our network (Supplementary Figure 1) led to a high correlation between these variables. Thus, ports with the highest connectivity to our target ports along the U.S. Gulf Coast are likely to play a disproportionate role in the dispersal of invasive mosquitoes to our target ports. Because the probability of unloading infested cargo from a given port diminishes with each unloading visit along a cargo ship’s route, and because accompanying invasive mosquitoes are most likely to survive and disperse given shorter travel times [22], we assumed (and modeled because first order Markov models are inherently weighted by distance) that ports most immediately visited by ships prior to arrival in US Gulf ports pose the greatest relative risk for importation of *Ae. aegypti* and *Ae. albopictus*.

While most ports on the US Gulf Coast have relatively little immediate connection to ports outside the Gulf Coast region, the high level of connectivity among several US Gulf ports (Supplementary Tables 1 and 2) may provide a vehicle for dispersal of invasive species into ports with less outside connectivity. Because *Ae. aegypti* and *Ae. albopictus* are so widely distributed among port cities, and especially those connected to ports in the US Gulf Coast, implementation of origin-specific screening is unlikely to lead to increased efficiency in halting the dispersal of these species into the US Gulf Coast region. Instead, preventing mosquitoes from entering the US Gulf Coast network seems particularly critical. Since Houston serves as a hub for vessels entering the US Gulf Coast network, implementation of an early alert and rapid response system for screening ships entering the Port of Houston could disproportionately reduce the risk of maritime dispersal of invasive species, including *Ae. aegypti* and *Ae. albopictus*. The findings from the agent-based model support the screening vessels upon arrival of destination as a strong intervention to reduce establishment of mosquito populations. Combined with the Markov model findings, mosquito removal of container cargo upon arrival to the Port of Houston could serve as an effective strategy at reducing invasive mosquito populations in the US Gulf Coast.

While our results accurately reflect the movements of all fully cellular container ships that arrived in the seven target ports along the US Gulf Coast, a number of potential routes of dispersal and potential vectors for dispersal were not considered in our study. First, our data did not include information on the movements of non-containerized cargo along the GSN. However, container ships are often considered to be better than non-containerized cargo ship as vectors for the dispersal of terrestrial invasive species because containers are rarely, if ever, opened and examined between destinations [43]. In addition, it is likely that our agent-based model conclusions, namely that screening at final destinations for mosquito infestations are the best way to prevent new invasions, holds true for smaller ships as these ships also visit ports within the *Aedes* spp. ranges.

Our models contained several assumptions and generalizations necessitated by data availability and the general lack of knowledge regarding transport of mosquitoes in cargo. Specifically, data collected by an automatic identification system does not include information on the number of containers or the type of cargo carried by each ship, so we assumed that each ship had the same capacity for infestation and transmission. Furthermore, these records do not include information on whether cargo was loaded or unloaded at each port. Some port visits are made for purposes of refueling, and involve no transfer of cargo to or from the vessel [44]. As a result, we assumed a constant probability for transmission from one port where a mosquito occurred to the next port, while in reality this probability is certainly heterogenous. This assumption was also present in our agent-based model, as was the feasibility of mosquito removal and screening of containerized cargo. More detailed information on containers and cargo, as well as quantification of mosquito infestation of these cargo, would dramatically improve our model and provide more insight into paths utilized by *Ae. aegypti* and *Ae. albopictus* for dispersal.

### Conclusions

This study represents the first pathway-based analysis of dispersal by *Ae. aegypti* and *Ae. albopictus* into and among major ports on the US Gulf Coast via the global shipping network. These mosquitoes, which are the primary vectors of numerous arboviruses that affect human an animal health [13–15], are also some of the most invasive insects on earth [9, 19]. Understanding long-distance dispersal of these species via maritime trade allows us to concentrate biosecurity and vector control efforts, thereby increasing management efficiency, and may allow us to better understand gene flow and patterns of population genetics and phenotypic traits that are important for mosquito control and public health [45]. We also show that port-based detection and eradication of potential mosquito invaders can substantially reduce the risks of martime-based invasion. A number of highly invasive and medically important mosquitoes, including *Anopheles stephensi* (Liston), *Ae*. (Hulecoeteomyia) *koreicus* and *Ae*. (Finlaya) *japonicus japonicus* (Theobald), are currently expanding their global ranges both over land and through long-distance dispersal via the GSN [46–48]. By understanding vector dispersal and its downstream effects, we may better understand and prevent outbreaks of vector-borne pathogens.

## Methods

We obtained data from Informa (formerly Lloyd’s Maritime Intelligence Unit; Informa, London, UK) detailing every fully cellular container ship that arrived at a major US port on the Gulf of Mexico in 2012 using automatic identification system transponders. We used these data and pathway-based, first-order Markov models to determine which ports along the US Gulf Coast were at the highest risk for importation of *Ae. aegypti* and *Ae. albopictus* along with container cargo shipped via maritime trade routes during 2012. Given that a ship loads and unloads cargo with each stop, our models also assume that some potential exists for infestation of the ship by mosquitoes at each stop at a port occupied by these species. These models therefore assume that some transmission potential exists between each port occupied by these species, and all ports visited subsequently. Thus, given a route A-B-C-D, where point D is the final port of call in the Gulf of Mexico and B is a port where at least one species of mosquito is present, we assume some potential for transmission from B to C and then from C to D. Because there is also some probability of cargo containing the mosquitoes to be unloaded at each port, we considered all points on a route together, running from *i* to *j*. This information was then used to assemble a database of routes *i* to *j* and the number of trips made by vessels along these routes.

Each route, *i* to *j*, had an associated number of stops *ij*. Each port occupied by either *Ae. aegypti* or *Ae. albopictus* was assigned constant transmission potential (*λ*) which was used to calculate the potential for importation (*P*_*ij*_) of each *Ae. aegypti* and *Ae. albopictus* into each one of our seven target ports. We then estimated the total relative likelihood of arrival by each species into each target port (*φ*_*j*_) by summing *P*_*ij*_ for all trips into each target port. Finally, we evaluated our model parameterization by generating multiple values for λ, and then generating a correlation matrix for *φ*_*ij*_ using Spearman rank correlation coefficients; models were robust to changes in parameterization (*r*_*s*_ > 0.964).

In the agent-based model, shipping containers aboard maritime ships were treated as agents, where mosquitoes found in these vessels could be moved between ports. Each container was assumed to start its journey with enough mosquitoes that mosquito establishment at new ports was theoretically possible. Containers moved between ports (varied from 1 to 10 stops) and these containers could be moved to shore at any given port with variable probabilities (ranged from 0 to 100% chance). Once moved to shore, mosquitoes had a set probability of establishing a population in the new location, ranging from 5 to 100%. When containers were not moved to shore, there was a 0 to 100% chance the container was transferred to a different cargo ship.

Under either condition, the mosquito population inside the container had an ∼90% chance of surviving at sufficient numbers to support a different establishment opportunity (individual survival probability determined as a random draw from a normal distribution with a mean of 0.9 and standard deviation of 0.035) to the next port. We assumed that the probability of a population establishing averaged 75% (agent survival probability ranged from ∼50-100% and was determined as a random draw from a normal distribution with mean of 0.75 and standard deviation of 0.09), which occurred just after containers were moved to shore. To combat the establishment of new mosquito vectors, we enacted port procedures for removing mosquitoes from shipping containers. Specifically, containers were checked upon arrival to shore or transfer between ships, with a probability of removing mosquitoes ranging from 0 to 100% for each event (Increment values for all probability ranges tested for each variable can be found in Supplement Table 3).

## Acknowledgements

We thank members of the Willoughby and Zohdy labs for constructive comments on previous versions of this manuscript. Funding for BAM was provided through a CDC-RFA-CK14-1401PPHF award and a USDA Young Investigator Research Award to SZ. This work was also partially supported by the USDA National Institute of Food and Agriculture, Hatch project 1025651 to JRW.

## Author contributions

JRW, BAM, and SZ conceived and designed the study. BAM and SZ collected the data and JRW, BAM, and JA performed analyses. JRW, BAM, JA, TDS, CAL, and SZ contributed to interpretation of the data as well as writing and editing the paper.

## Data accessibility

Ship movement and port of call data included in Supplementary Table 4. All agent based model scripts and analysis code is available via GitHub: https://github.com/jwillou/maritime_transport.

## Disclaimer

The findings and conclusions in this report are those of the authors and do not necessarily represent the official position of the Centers for Disease Control and Prevention.

## Notes

### Competing Interest Statement

The authors have declared no competing interest.

## References

1. Glaesser D, Kester J, Paulose H, et al (2017) Global travel patterns: an overview. J Travel Med 24:. https://doi.org/10.1093/jtm/tax007

2. Sardain A, Sardain E, Leung B (2019) Global forecasts of shipping traffic and biological invasions to 2050. Nat Sustain 2:274–282

3. Hulme PE (2009) Trade, transport and trouble: managing invasive species pathways in an era of globalization. J Appl Ecol 46:10–18

4. Banks NC, Paini DR, Bayliss KL, Hodda M (2015) The role of global trade and transport network topology in the human-mediated dispersal of alien species. Ecol Lett 18:188–199

5. Keller RP, Drake JM, Drew MB, Lodge DM (2011) Linking environmental conditions and ship movements to estimate invasive species transport across the global shipping network. Divers Distrib 17:93–102

6. Drake JM, Lodge DM (2004) Global hot spots of biological invasions: evaluating options for ballast-water management. Proc Biol Sci 271:575–580

7. Sylvester F, MacIsaac HJ (2010) Is vessel hull fouling an invasion threat to the Great Lakes? Divers Distrib 16:132–143

8. Paini DR, Yemshanov D (2012) Modelling the arrival of invasive organisms via the international marine shipping network: a Khapra beetle study. PLoS One 7:e44589

9. Bonizzoni M, Gasperi G, Chen X, James AA (2013) The invasive mosquito species Aedes albopictus: current knowledge and future perspectives. Trends Parasitol 29:460–468

10. Powell JR, Tabachnick WJ (2013) History of domestication and spread of Aedes aegypti--a review. Mem Inst Oswaldo Cruz 108 Suppl 1:11–17

11. Kampen H, Werner D (2014) Out of the bush: the Asian bush mosquito Aedes japonicus japonicus (Theobald, 1901) (Diptera, Culicidae) becomes invasive. Parasit Vectors 7:59

12. Gratz NG (2004) Critical review of the vector status of Aedes albopictus. Med Vet Entomol 18:215–227

13. Paupy C, Delatte H, Bagny L, et al (2009) Aedes albopictus, an arbovirus vector: from the darkness to the light. Microbes Infect 11:1177–1185

14. McKenzie BA, Wilson AE, Zohdy S (2019) Aedes albopictus is a competent vector of Zika virus: A meta-analysis. PLoS One 14:e0216794

15. Souza-Neto JA, Powell JR, Bonizzoni M (2019) Aedes aegypti vector competence studies: A review. Infect Genet Evol 67:191–209

16. Guerra CA, Reiner RC Jr, Perkins TA, et al (2014) A global assembly of adult female mosquito mark-release-recapture data to inform the control of mosquito-borne pathogens. Parasit Vectors 7:276

17. Benedict MQ, Levine RS, Hawley WA, Lounibos LP (2007) Spread of the tiger: global risk of invasion by the mosquito Aedes albopictus. Vector Borne Zoonotic Dis 7:76–85

18. Kraemer MUG, Sinka ME, Duda KA, et al (2015) The global distribution of the arbovirus vectors Aedes aegypti and Ae. albopictus. Elife 4:e08347

19. Kraemer MUG, Reiner RC Jr, Brady OJ, et al (2019) Past and future spread of the arbovirus vectors Aedes aegypti and Aedes albopictus. Nat Microbiol 4:854–863

20. Hawley WA (1988) The biology of Aedes albopictus. J Am Mosq Control Assoc Suppl 1:1–39

21. Braks MAH, Honório NA, Lounibos LP, et al (2004) Interspecific Competition Between Two Invasive Species of Container Mosquitoes, Aedes aegypti and Aedes albopictus (Diptera: Culicidae), in Brazil. Ann Entomol Soc Am 97:130–139

22. Brown HE, Smith C, Lashway S (2017) Influence of the length of storage on Aedes aegypti (Diptera: Culicidae) egg viability. J Med Entomol 54:489–491

23. Reiter P, Sprenger D (1987) The used tire trade: a mechanism for the worldwide dispersal of container breeding mosquitoes. J Am Mosq Control Assoc 3:494–501

24. Pliego Pliego E, Velázquez-Castro J, Eichhorn MP, Fraguela Collar A (2018) Increased efficiency in the second-hand tire trade provides opportunity for dengue control. J Theor Biol 437:126–136

25. Monaghan AJ, Sampson KM, Steinhoff DF, et al (2018) The potential impacts of 21st century climatic and population changes on human exposure to the virus vector mosquito Aedes aegypti. Clim Change 146:487–500

26. Liu B, Gao X, Ma J, et al (2019) Modeling the present and future distribution of arbovirus vectors Aedes aegypti and Aedes albopictus under climate change scenarios in Mainland China. Sci Total Environ 664:203–214

27. Gubler DJ (2002) The global emergence/resurgence of arboviral diseases as public health problems. Arch Med Res 33:330–342

28. Zohdy S, Morse WC, Mathias D, et al (2018) Detection of Aedes (Stegomyia) aegypti (Diptera: Culicidae) populations in southern Alabama following a 26-yr absence and public perceptions of the threat of Zika virus. J Med Entomol. https://doi.org/10.1093/jme/tjy050

29. Colunga-Garcia M, Haack R, Magarey R, Borchert D (2013) Understanding trade pathways to target biosecurity surveillance. NeoBiota 18:103–118

30. Simpson A, Jarnevich C, Madsen J, et al (2009) Invasive species information networks: collaboration at multiple scales for prevention, early detection, and rapid response to invasive alien species. Biodiversity (Nepean) 10:5–13

31. Monaghan AJ, Morin CW, Steinhoff DF, et al (2016) On the seasonal occurrence and abundance of the Zika virus vector mosquito Aedes aegypti in the contiguous United States. PLoS Curr. https://doi.org/10.1371/currents.outbreaks.50dfc7f46798675fc63e7d7da563da76

32. Tesla B, Demakovsky LR, Mordecai EA, et al (2018) Temperature drives Zika virus transmission: evidence from empirical and mathematical models. Proc Biol Sci 285:20180795

33. Graham AS, Pruszynski CA, Hribar LJ, et al (2011) Mosquito-associated dengue virus, Key West, Florida, USA, 2010. Emerg Infect Dis 17:2074–2075

34. Kendrick K, Stanek D, Blackmore C, Centers for Disease Control and Prevention (CDC) (2014) Notes from the field: Transmission of chikungunya virus in the continental United States--Florida, 2014. MMWR Morb Mortal Wkly Rep 63:1137

35. Likos A, Griffin I, Bingham AM, et al (2016) Local mosquito-borne transmission of Zika virus - Miami-Dade and Broward counties, Florida, June-August 2016. MMWR Morb Mortal Wkly Rep 65:1032–1038

36. Musso D, Rodriguez-Morales AJ, Levi JE, et al (2018) Unexpected outbreaks of arbovirus infections: lessons learned from the Pacific and tropical America. Lancet Infect Dis 18:e355–e361

37. Chouin-Carneiro T, Vega-Rua A, Vazeille M, et al (2016) Differential Susceptibilities of Aedes aegypti and Aedes albopictus from the Americas to Zika Virus. PLoS Negl Trop Dis 10:e0004543

38. Azar SR, Roundy CM, Rossi SL, et al (2017) Differential Vector Competency of Aedes albopictus Populations from the Americas for Zika Virus. Am J Trop Med Hyg 97:330–339

39. Ciota AT, Bialosuknia SM, Zink SD, et al (2017) Effects of Zika virus strain and Aedes mosquito species on vector competence. Emerg Infect Dis 23:1110–1117

40. Ramos MM, Mohammed H, Zielinski-Gutierrez E, et al (2008) Epidemic dengue and dengue hemorrhagic fever at the Texas-Mexico border: results of a household-based seroepidemiologic survey, December 2005. Am J Trop Med Hyg 78:364–369

41. Alto BW, Wiggins K, Eastmond B, et al (2017) Transmission risk of two chikungunya lineages by invasive mosquito vectors from Florida and the Dominican Republic. PLoS Negl Trop Dis 11:e0005724

42. United States. Bureau of Transportation Statistics (2020) Port performance freight statistics program: Annual report to congress 2019

43. Derraik JGB (2004) Exotic mosquitoes in New Zealand: a review of species intercepted, their pathways and ports of entry. Aust N Z J Public Health 28:433–444

44. Ducruet C, Notteboom T (2012) The worldwide maritime network of container shipping: spatial structure and regional dynamics. Glob Netw (Oxf) 12:395–423

45. Manni M, Guglielmino CR, Scolari F, et al (2017) Genetic evidence for a worldwide chaotic dispersion pattern of the arbovirus vector, Aedes albopictus. PLoS Negl Trop Dis 11:e0005332

46. Medlock JM, Hansford KM, Schaffner F, et al (2012) A review of the invasive mosquitoes in Europe: ecology, public health risks, and control options. Vector Borne Zoonotic Dis 12:435–447

47. Kaufman MG, Fonseca DM (2014) Invasion biology of Aedes japonicus japonicus (Diptera: Culicidae). Annu Rev Entomol 59:31–49

48. Surendran SN, Sivabalakrishnan K, Sivasingham A, et al (2019) Anthropogenic factors driving recent range expansion of the malaria vector Anopheles stephensi. Front Public Health 7:53

