## Supplemental for "Mosquito invasion via the global shipping network is slowed in high-risk areas by on-shore and ship-board monitoring"

**Supplementary information**

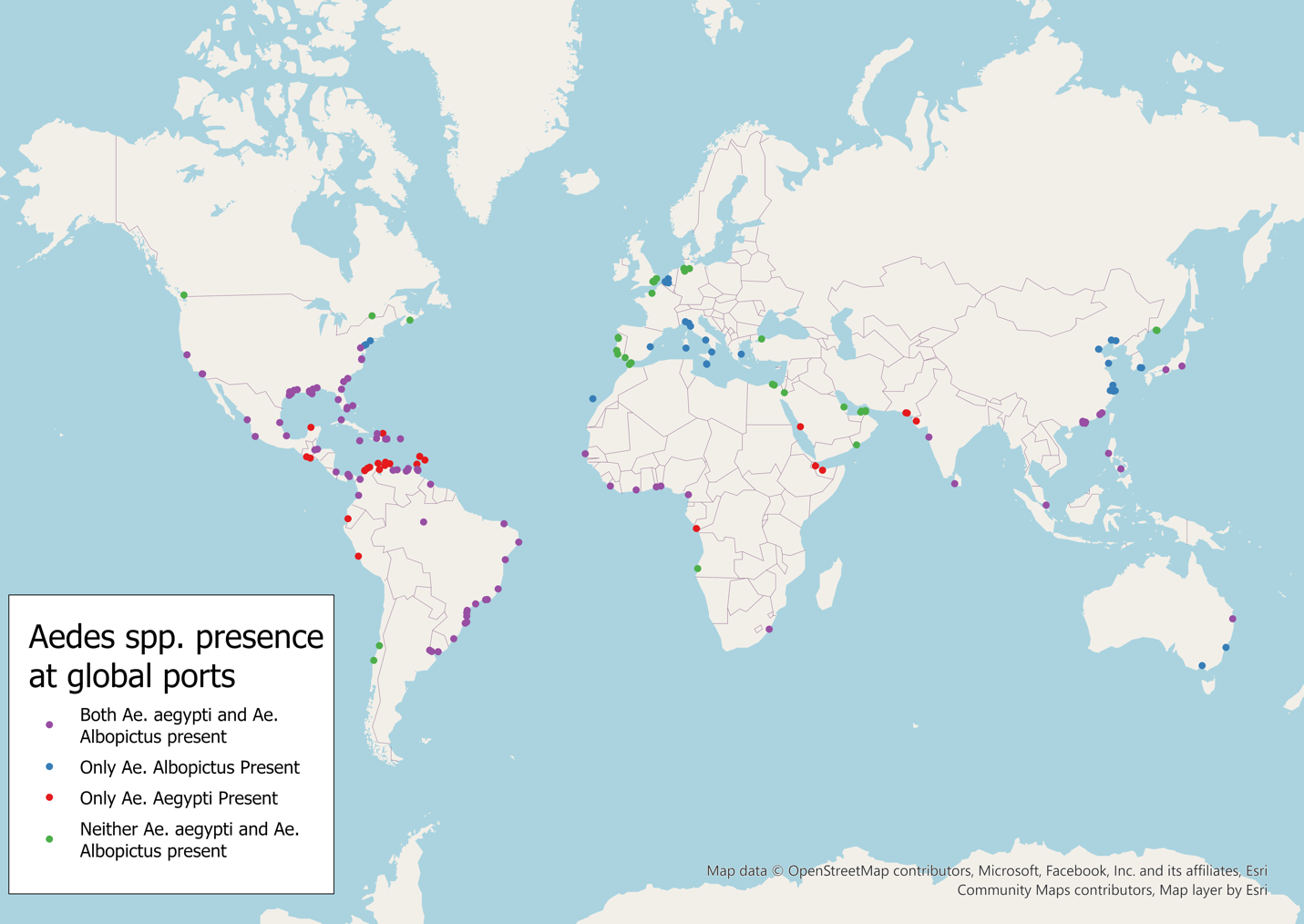

**Supplementary Figure 1.** Using *Aedes* distributions maps [18], we determined that only 39 (18.3%) of the 213 ports within our network (distributed across 69 countries) were free of *Ae. aegypti* and *Ae.albopictus* populations; 140 (65.7%) ports within our network hosted populations of *Ae. aegypti*, 148 (69.4%) hosted populations of *Ae. albopictus*, and 114 (53.5%) ports within our network hosted populations of both *Ae. aegypti* and *Ae. albopictus.*

**Supplementary Table 1.** Ports with the highest immediate connectivity to our seven target ports in the US Gulf States. Since nearly all maritime arrivals in the Gulf passed most recently through ports on the Atlantic seaboard, in the Caribbean, or in other ports on the Gulf of Mexico, all of which host populations of both *Ae. aegypti* and *Ae. albopictus*, mosquito populations from these ports must reasonably be assumed to be the most likely to arrive in target ports. Data represents arrivals by fully cellular container ships from January 1^st^ to December 31^st^, 2012.

| International Ports or U.S. Ports Outside Gulf of Mexico | Total trips to target ports | Trips to Houston | Trips to New Orleans | Trips to Mobile | Trips to Gulfport | Trips to Tampa |
| --- | --- | --- | --- | --- | --- | --- |
| Altamira, Mexico | 373 | 364 | 0 | 9 | 0 | 0 |
| Santo Tomás de Castilla, Guatemala | 160 | 130 | 29 | 0 | 1 | 0 |
| Puerto Cortes, Honduras | 105 | 3 | 5 | 0 | 96 | 0 |
| Savannah, Georgia, USA | 104 | 102 | 0 | 2 | 0 | 0 |
| Kingston, Jamaica | 102 | 17 | 18 | 18 | 0 | 49 |

**Supplementary Table 2.** High connectivity between ports on the US Gulf Coast implies high risk for movement of *Aedes* spp. mosquitoes between these cities. While Houston seems to play a role as a hub for international arrivals, New Orleans and Mobile receive a great number of shipments from domestic ports, including Houston. Data represents arrivals by fully cellular container ships from January 1^st^ to December 31^st^, 2012.

| U.S. Ports along Gulf of Mexico | Trips to Houston | Trips to New Orleans | Trips to Mobile | Trips to Gulfport | Trips to Freeport | Trips to Tampa | Trips to Galveston |
| --- | --- | --- | --- | --- | --- | --- | --- |
| Houston, TX | NA | 316 | 98 | 0 | 6 | 0 | 2 |
| New Orleans, LA | 89 | NA | 50 | 0 | 0 | 0 | 0 |
| Mobile, AL | 32 | 39 | NA | 0 | 0 | 7 | 0 |
| Gulfport, MS | 0 | 0 | 0 | NA | 0 | 0 | 0 |
| Freeport, TX | 0 | 6 | 0 | 0 | NA | 0 | 0 |
| Tampa, FL | 0 | 0 | 48 | 0 | 0 | NA | 0 |
| Galveston, TX | 2 | 0 | 0 | 0 | 0 | 0 | NA |

**Supplementary Table 3.** Description of parameters and the values of each parameter we tested in our agent-based models. All pairwise combinations of values (in total 784,080 sets) were tested and replicated 100 times. Default parameters values were used when isolating the effects of individual parameters on mosquito population establishment.

| **parameter** | **value range** | **increment** | **default** |
| --- | --- | --- | --- |
| probability that a container was moved from ship to shore | 0-100% | 0.1 | 50% |
| probability that a container was moved from ship to a different ship | 0-100% | 0.1 | 50% |
| probability of mosquitoes leaving an onshore container and establishing a viable population | 50-100% | 0.1 | 75% |
| number of stops on a ship route | 1-10 | 1 | 5% |
| probability of mosquitoes surviving the journey between ports | 80-100% | 0.1 | 90% |
| probability of detecting and removing mosquitoes from on-land container | 0-100% | 0.25 | 40% |
| probability of detecting and removing mosquitoes from ship-board container | 0-100% | 0.25 | 40% |

**Supplementary Table 4.** Maritime transport data from ship transponders and shore receivers used in these analyses. Columns include an individual ship identification number, a column noting the arrival port and a column noting the departure port.
